# Presynaptic GABAergic receptors modulate inhibitory synaptic feedback in the avian cochlear nucleus angularis

**DOI:** 10.1101/619783

**Authors:** Stefanie L. Eisenbach, Sara E. Soueidan, Katrina M. MacLeod

## Abstract

Inhibition plays multiple critical roles in the neural processing of sound. In the avian auditory brain stem, the cochlear nuclei receive their principal inhibitory feedback from the superior olivary nucleus (SON) in lieu of local inhibitory circuitry. In the timing pathway, GABAergic inhibitory feedback underlies gain control to enhance sound localization. In the cochlear nucleus angularis (NA), which processes intensity information, how the inhibitory feedback is integrated is not well understood. Using whole cell patch-clamp recordings in chick brain stem slices, we investigated the effects of GABA release on the inhibitory (presumed SON) and excitatory (8^th^ nerve) synaptic inputs onto NA neurons. Pharmacological activation of the metabotropic GABA_B_ receptors with baclofen profoundly suppressed both evoked excitatory and inhibitory postsynaptic currents (EPSCs and IPSCs). Baclofen similarly reduced the frequency of spontaneous IPSCs and EPSCs, but had no significant effect on the current kinetics or amplitudes, indicating a presynaptic locus of modulation. Trains of IPSCs showed substantial transient and sustained short-term synaptic facilitation. Baclofen application reduced the initial IPSC amplitude, but enhanced the relative facilitation over the train via changes in release probability. Comparable levels of GABA_B_ receptor mediated blockade also shifted short-term synaptic plasticity of EPSCs toward less depression. Evoked (but not spontaneous) release of GABA was sufficient to suppress basal release at inhibitory synapses in slices. Overall, the modulation of excitatory and inhibitory inputs of NA neurons via GABA_B_ receptor activation appears to parallel that in the timing pathway.

**New and Noteworthy:** Avian cochlear nucleus angularis (NA) neurons are responsible for encoding sound intensity and provide level information for gain control feedback via the superior olivary nucleus. This GABAergic inhibitory feedback was itself modulated in NA via presynaptic, metabotropic GABA_B_ receptor mediated suppression. Excitatory transmission was modulated by the same receptors, suggesting parallel homosynaptic and heterosynaptic mechanisms in both cochlear nuclei.

## Introduction

Inhibition plays a critical role in auditory processing, contributing in various ways to sound level gain control (Peña et al., 1996), sculpting frequency tuning (Köppl and Carr, 2003; Fukui et al., 2010), and enhancing temporal processing (Funabiki et al., 1998; Yamada et al., 2013). In the avian system, nearly all cochlear nucleus neurons provide feedforward excitatory projections to the ascending pathways, while inhibitory inputs principally arise from an anatomically separate brain stem region, the superior olivary nucleus (SON) (Funabiki et al., 1998; Yang et al., 1999; Lu and Trussell, 2000; Monsivais et al., 2000; Lu and Trussell, 2001). The SON provides ipsilateral feedback inhibition to both divisions of the cochlear nucleus, nucleus angularis (NA) and nucleus magnocellularis (NM), as well as to nucleus laminaris (NL) (Burger et al., 2005a; Nishino et al., 2008). This inhibitory feedback loop to NM and NL has been proposed to enact an intensity-dependent gain control that enhances coincidence detection for interaural time difference. While NA provides a source of sound level information to the SON (Lachica et al., 1994; Yang et al., 1999), how the reciprocal inhibitory feedback is integrated into sound processing in NA is not yet clear.

The mechanisms underlying inhibitory control of the timing nuclei involve both presynaptic and postsynaptic regulation. Inhibitory synaptic terminals in these nuclei release the neurotransmitter γ–aminobutyric acid (GABA), along with glycine (Kuo et al., 2009). Postsynaptic inhibitory responses are principally GABA_A_ receptor mediated in NM (Tang et al., 2009; Coleman et al., 2011; Tang and Lu, 2012a; Fischl and Burger, 2014), while responses in NA are mediated by both GABA_A_ and glycine receptors (Kuo et al. 2009). Presynaptic inhibition is mediated via metabotropic GABA (GABA_B_) receptors in NM: pharmacological activation of GABA_B_ receptors has a strong presynaptic suppressive effect on glutamatergic inputs to NM neurons (Brenowitz et al., 1998; Brenowitz and Trussell, 2001a; 2001b). Activation of GABA_B_ receptors also suppressed inhibitory inputs onto NM neurons, as well as onto NL neurons (Tang et al., 2009), suggesting the inhibitory terminals were self-regulatory via homosynaptic feedback.

Less is known about the mechanisms and functional effects of presynaptic inhibition in nucleus angularis. Immunohistochemical and electron microscopy studies showed GABA_B_ receptors were also expressed in NA (Burger et al., 2005b). Like in NM, a previous study has shown that GABA_B_ agonists had a suppressive effect on excitatory synaptic inputs into NA via presynaptic mechanisms (Shi and Lu, 2017). In this study, we examined the effects of activation of GABA_B_ mechanisms on the inhibitory inputs to NA neurons using in vitro whole cell patch clamp electrophysiology in the brain stem slice preparation. The results provide evidence for GABAergic presynaptic modulation of these putative SON inputs to NA, as well as similar modulation of excitatory (presumptive nerve) input. These data suggest a common mechanism for homosynaptic regulation was present at the inhibitory terminals at all three descending targets of the SON.

## Methods

### Chick brain stem slice preparation

Embryonic day 17-18 chicken embryos were anesthetized and rapidly decapitated in accordance with the University of Maryland Institutional Animal Care and Use Committee guidelines. The brain stem containing the cochlear nuclei was dissected out of the skull and submerged in low sodium artificial cerebral spinal fluid (ACSF) (in mM: 97.5 NaCl, 3 KCl, 2.5 MgCl_2_, 26 NaHCO_3_, 2 CaCl_2_, 1.25 NaH_2_PO_4_, 10 dextrose, 3 HEPES, 57.5 sucrose) that was being continuously oxygenated with 95% O_2_- 5% CO_2_. Transverse brain stem slices 250 μm thick were cut using a vibrating tissue slicer (Leica Microsystems, Wetzlar, Germany), then incubated in oxygenated normal ACSF (in mM: 130 NaCl, 3 KCl, 2 MgCl_2_, 26 NaHCO_3_, 2 CaCl_2_, 1.25 NaH_2_PO_4_, 10 dextrose, 3 HEPES) heated to 34°C for 30 minutes, then held at room temperature.

### Patch-clamp electrophysiology

For recordings, slices were transferred to a submersion recording chamber continuously perfused with oxygenated ACSF (1–2 ml min^−1^) and heated with a Warner TC 324B inline heating device (Warner Instr., Hamden, CT) to 30 ± 0.2°C. Whole cell patch-clamp recordings were made from visually-identified nucleus angularis neurons using IR/DIC (infrared/differential interference contrast) video microscopy. Electrophysiological recordings were made with a Multiclamp 700B patch clamp amplifier (Molecular Devices, Sunnyvale, CA) in voltage-clamp mode. Initial micropipette resistances were 4.5–7 MΩ. The series resistance was corrected by 60–90% and the junction potentials were left uncorrected. Stimulation and recordings were controlled by a PC running custom software in IGOR Pro (Wavemetrics, Lake Oswego, OR). Two intracellular solutions were used: excitatory (AMPA-receptor mediated) synaptic currents were measured with a cesium-based intracellular voltage-clamp solution (in mM: 70 cesium sulfate, 5 QX-314 (lidocaine), 1 MgCl_2_, 1 Na_2_ATP, 0.3 Na_2_GTP, 10 phosphocreatine, 4 NaCl, 10 HEPES ((4-(2-hydroxyethyl)-1-piperazineethanesulfonic acid), 5 BAPTA (1,2-bis(o-aminophenoxy)ethane-N,N,N’,N’-tetraacetic acid), 32 sucrose and 0.2% biocytin). Inhibitory (mixed GABA/glycine receptor mediated) synaptic currents were recorded with a similar solution with cesium chloride to achieve a chloride reversal potential was close to zero mV (in mM: 130 cesium chloride, 5 QX-314, 1 MgCl_2_, 1 Na_2_ATP, 0.3 Na_2_GTP, 5 phosphocreatine, 10 HEPES, 5 BAPTA, and 0.2% biocytin). IPSCs that were recorded under voltage clamp near resting potential were therefore measured as inward currents. To eliminate GABA/glycine inhibitory currents for EPSC recordings, gabazine (20 μM) and strychnine, (3 μM) were included in the bath ACSF. Alternatively, to eliminate glutamatergic synaptic currents for IPSC recordings DNQX (10 μM) and APV (25 μM) were included. GABA_B_ receptor agonist baclofen (20-50 μM, Sigma), antagonist CGP 52432 (10-20 μM, Tocris Cookson) and GABA transporter inhibitor NNC-771 (20-80 μM, Tocris Cookson) were also bath applied. Spontaneous events were recorded in the absence of tetrodotoxin as there is no evidence for spontaneous firing activity in the cochlear nucleus in slice under our recording conditions and a previous study showed no differences between spontaneous excitatory synaptic events and so-called “miniature events” recorded in the presence of tetrodotoxin (MacLeod and Carr, 2005). Spontaneous IPSC events measured using 20 μM (n=4) and 80 μM (n=4) NNC 771 were pooled as experiments showed no differences in event properties or drug effect between concentrations. Spontaneous sIPSC events measured with 10 μM (n=6) and 20 μM (n=2) CGP 52432 were also pooled.

Excitatory synaptic currents were evoked with a tungsten monopolar electrode placed amongst the 8^th^ (auditory) nerve fibers tracts as they enter the nucleus angularis at the medial margin. Inhibitory syanptic currents were more difficult to isolate, but generally were stimulated from within the nucleus boundary, most often 100-200 μm medially or ventrally distant relative to the recording site. Biphasic stimulus waveforms passed through an analog, constant-current stimulus isolation unit (World Precision Instruments, Sarasota, FL). Stimulus artifacts were digitally removed for clarity in figure presentations.

### Analysis and statistics

Spontaneous [E/I]PSC kinetics were analyzed for recordings in which there were sufficient numbers of events to test in both control and drug conditions (>30 events per condition). Paired pulse ratio (PPR) was the second PSC amplitude relative to the initial amplitude; the steady state ratio (SSR) was the average of PSC 8 through 10 of the train relative to the initial PSC amplitude. Single or double exponential fits were performed using built in curve fitting in Igor Pro v.8. Weighted time constants for double exponential fits were according to 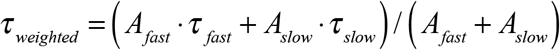 and fast time constant percentage was calculated as 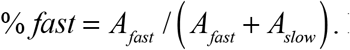. Distributions were tested for differences by two-sample Kolomogorov-Smirnov test with a signficance threshold of p<0.05. Summary statistics were calculated using Microsoft Excel. Values reported in the text are mean±s.d. unless otherwise indicated. Hypothesis testing used Student’s paired *t* Test unless otherwise indicated. Short-term plasticity during trains were analyzed with 2-way ANOVA with two within subject factors of drug condition and stimulus number, and post hoc pairwise Student’s *t* Test with Bonferroni correction for multiple comparisons. Statistical analysis was performed using Microsoft Excel or the statistical software *R* (Version 0.98.1103 RStudio, Inc.) on a MacBook Pro.

## Results

### Activation of metabotropic GABA receptors suppresses inhibitory and excitatory neurotransmitter release

To investigate inhibitory modulation of synaptic transmission in the avian cochlear nucleus angularis, the specific GABA_B_ receptor agonist baclofen was applied during recordings of inhibitory postsynaptic currents (IPSCs) in acute brain stem slices. Synaptic currents were evoked by extracellular electrical stimulation and measured under whole-cell voltage clamp mode. Baclofen (20 μM) reliably reduced the peak amplitudes of pharmacologically isolated evoked IPSCs (Fig. 1A). On average, baclofen reduced IPSC amplitudes to 10.5% of control levels (control: −0.38±0.18 nA, baclofen: −0.041±0.043 nA, n=12, p=2.4×10^−5^, Student’s paired *t* Test; Fig. 1B). The baclofen effect on IPSC amplitudes was partially and inconsistently reversed with washout of the drug (blue shaded region in Fig. 1A), recovering to 52% of control levels on average (control: −0.37±0.19 nA, washout: −0.20±0.16 nA, n=10, p=0.0032).

**Figure 1.**
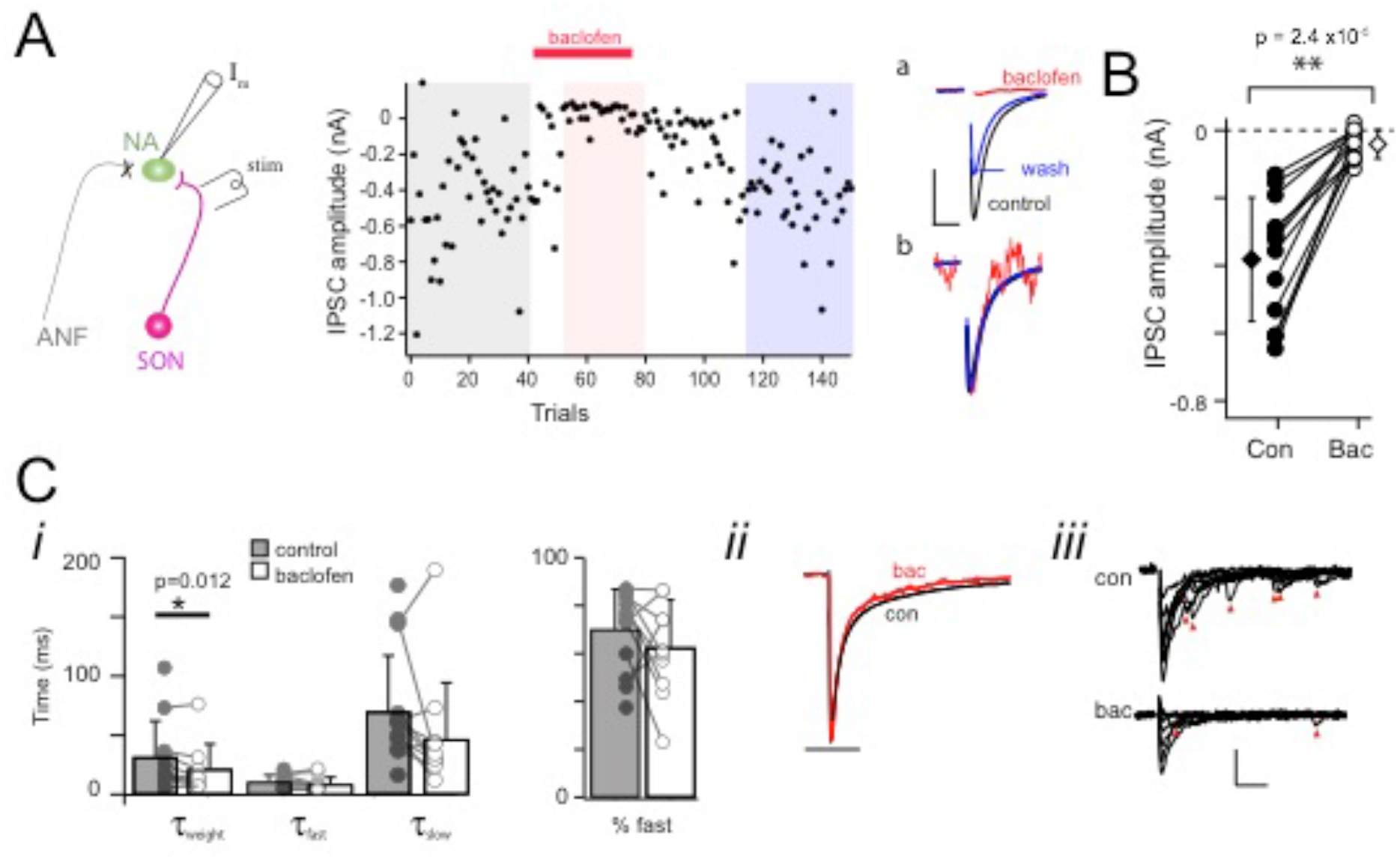
Baclofen suppressed evoked inhibitory postsynaptic currents onto neurons in the cochlear nucleus angularis (NA). A) Schematic of recording arrangement (left). Time plot of amplitudes of evoked single IPSCs elicited by stimulating electrode at a low rate (0.1 Hz) before, during, and after bath application of baclofen (20 μM)(middle). Right: average IPSC traces in control (black trace), baclofen (red) and during partial washout (blue) overlaid (a) or normalized and overlaid (b). Traces correspond to averages of responses in gray, red and blue shaded regions, respectively, in time plot. ANF, auditory nerve fiber, SON, superior olivary nucleus. B) Summary data of IPSC absolute amplitude (n=12). Baclofen reduced the IPSCs by >90%. ** Student’s paired *t* Test, p=2.4×10^−5^. C) Kinetics analysis of the decay of the IPSCs fit with double exponential showed a slight decrease in time constants *τ_fast_* and *τ_slow_*, with significant effect on weighted time constant *τ_weight_* (*i*). Grand average of all normalized IPSCs before (control) and after baclofen (*ii*). Individual traces from one recording showed numerous asynchronous events (red arrowheads) that likely contribute to the slower decay and were largely absent in baclofen (*iii*). Exponential fits of grand averages: *τ_fast_* =8.4 ms, *τ_slow_* =93 ms, % *fast* = 68% in control; *τ_fast_* =6.8 ms, *τ_slow_* =63.0 ms, % *fast* = 72% in baclofen. Horizontal scale bars: 10 ms (Aa), 50 ms (Cii), 20 ms (Ciii). Vertical scale bars: 200pA (Aa), 100 pA (C*iii*).

Baclofen had little effect on the time course of the evoked currents (Fig. 1Ab). Most evoked IPSCs could be best fit with a double exponential with a fast and slow time constant in control which marginally decreased with baclofen (control: *τ_fast_* =10.3±5.0 ms, *τ_slow_* =66.2±37.7 ms; baclofen: *τ_fast_* =8.3±6.3 ms, *τ_slow_* =46.6±47.9 ms; n=10, no significant difference, p=0.12 and p=0.14, respectively; Fig. 1Ci). A small but significant decrease in the weighted time constant of the current decay suggested a slight speeding up of the overall current (control: *τ_weight_* 26.4±19.4 ms, baclofen: *τ_weight_* =21.4±21.3 ms, n=10, p=0.012, Student’s paired *t* Test). This effect may be due to the reduction in asynchronous events that could be seen in some recordings along with a decrease in the synchronous IPSC (Fig. 1Ciii).

We observed similar levels of GABA_B_ receptor driven suppression on evoked EPSCs, the putative auditory nerve inputs (Fig. 2A). Baclofen (20 μM) reliably suppressed the mean EPSC peak amplitude to 8.4% of control levels (control: −0.48±0.32 nA; baclofen: 0.040±0.048 nA, n=7, p=0.011; Fig. 2B). Washout of the baclofen effect on EPSCs was slow and inconsistent, only measurable in a subset of neurons (control: −0.57±0.37, washout: −0.71±0.85, n=4, no significant difference, p=0.60)(Fig. 2A). Baclofen had no significant effect on the time course of evoked EPSCs fit with a single exponential curve (control: *τ* = 2.7±1.7 ms, baclofen: *τ* = 2.6±0.8 ms n=6, no significant difference, p=0.70; Fig. 2C).

**Figure 2.**
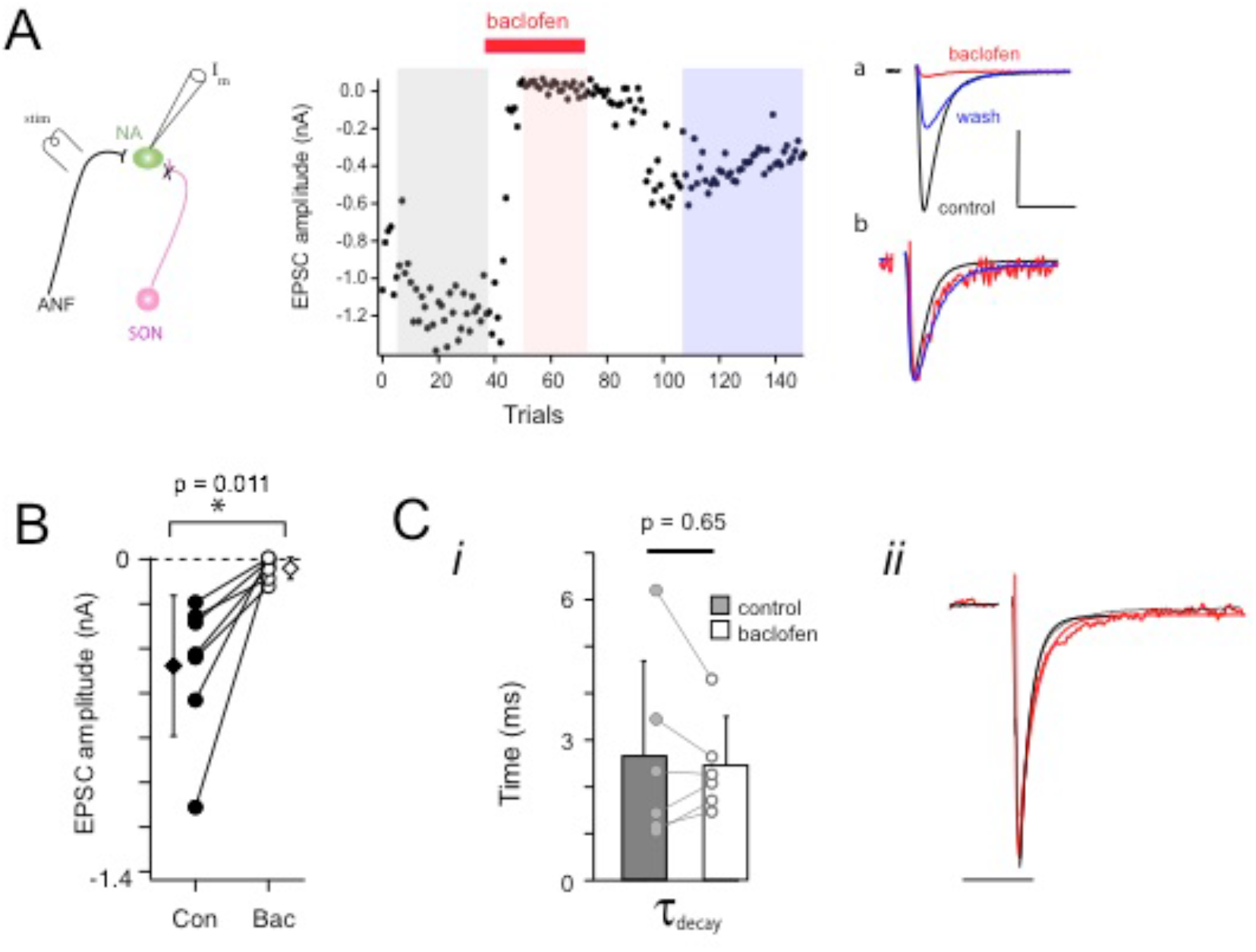
Baclofen suppressed evoked excitatory postsynaptic currents onto neurons in the cochlear nucleus angularis. A) Schematic of recording arrangement (left). Time plot of amplitudes of evoked single EPSCs evoked at a low rate (0.1Hz) before, during, and after bath application of baclofen (20 μM)(middle). Right: average EPSC traces in control (black trace), baclofen (red) and during partial washout (blue) overlaid (a) or normalized and overlaid (b). Traces correspond to averages of responses in gray, red and blue shaded regions, respectively, in time plot. ANF, auditory nerve fiber, SON, superior olivary nucleus. B) Summary data of EPSC absolute amplitude (n=7). Baclofen also reduced the EPSCs by >90%. * Student’s paired *t* Test, p=0.011. C) Kinetics analysis of the decay of the EPSCs fit with single exponential showed no significant effect (*i*). Grand average of all normalized EPSCs before (control) and after baclofen (*ii*). Exponential fits of grand averages: *τ* =1.6 ms in control; *τ* =2.5 ms in baclofen. Horizontal scale bars: 5 ms (Aa), 10 ms (Cii). Vertical scale bar: 600pA (Aa).

For both excitatory and inhibitory synapses, the baclofen effect was blocked by co-application of the specific GABA_B_ receptor antagonist, CGP52432 (10 μM). Blockade was complete for evoked IPSCs with no signficant difference between control IPSC amplitudes IPSCs in presence of agonist and antagonist together (control: −0.53±0.23 nA; baclofen+CGP: −0.53±0.11 nA, n=4, p=0.93). The same concentration of CGP52432 also substantialy blocked the baclofen suppression of the evoked EPSCs, retaining 74.4% of control amplitudes (control: −0.90±0.54 nA; baclofen+CGP: −0.64±0.40 nA, n=5, p=0.055).

### Presynaptic locus of suppression

To determine whether baclofen was acting presynaptically, we examined the effect of the drug on pharmacologically isolated spontaneous synaptic currents under voltage clamp. Baclofen application significantly decreased the frequency of both inhibitory and excitatory spontaneous PSCs (sIPSCs and sEPSCs, respectively)(Fig. 3Ai, Bi). Cumulative histograms of the interevent intervals for sIPSCs showed significant shifts to longer intervals (N=400 control events, N=397 baclofen events, p=1.12×10^−10^, Kolmogorov-Smirnov test, Fig.3Aii). The average spontaneous event rate for sIPSCs was reduced by >70% when baclofen was bath applied (control: 2.1±1.7 Hz, baclofen: 0.60±0.76 Hz, n=20, p=1.9×10^−6^, Wilcoxon rank sum test)(Fig. 3Ci).

**Figure 3.**
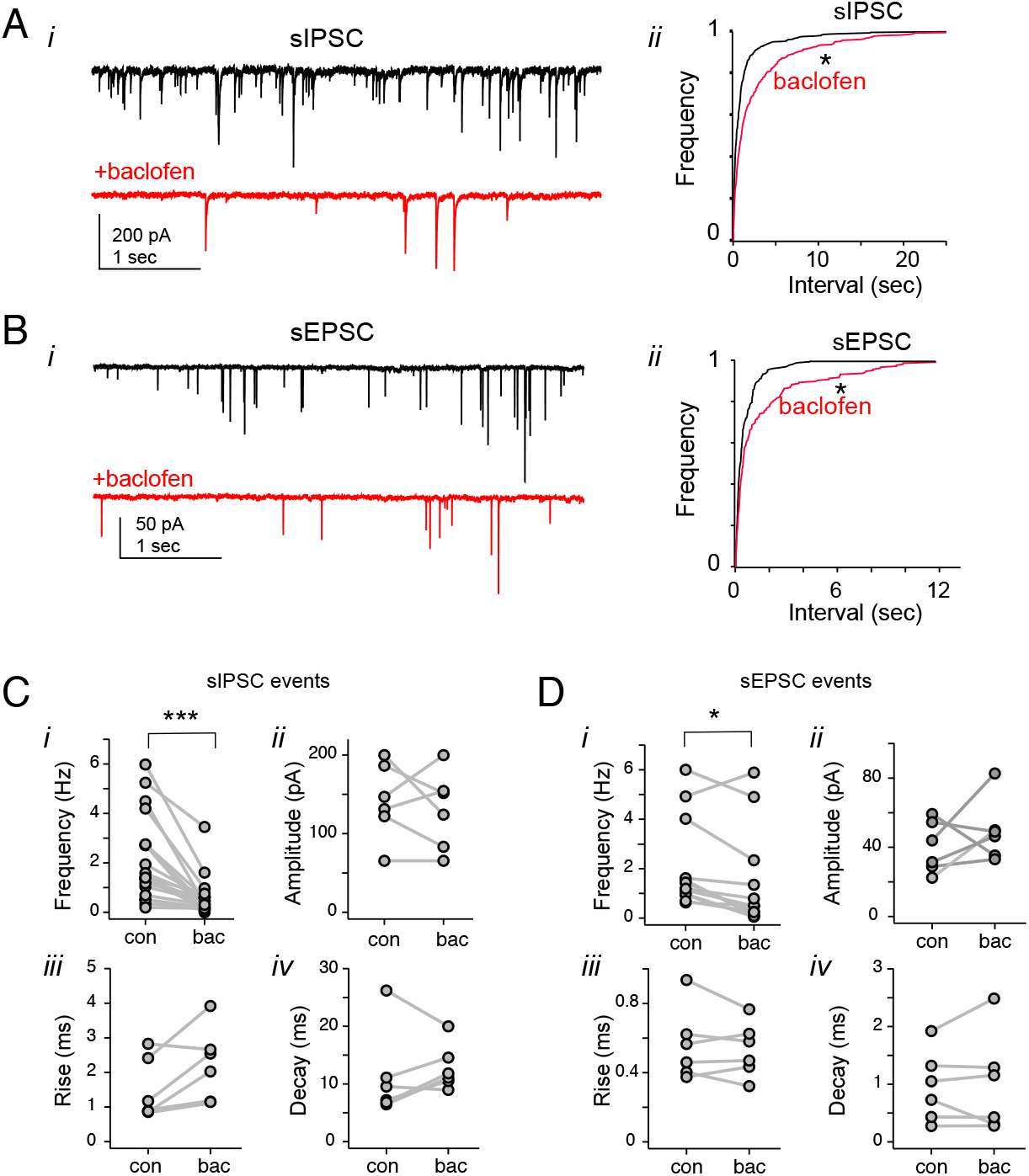
Baclofen effects on spontaneous IPSC and EPSC events.Baclofen application (50 μM) reduced the frequency of spontaneous IPSC events (A) and spontaneous EPSC events (B). Current traces (i) and cumulative histograms of PSC intervent intervals (ii) before (black) and after application of baclofen (red). Baclofen shifted the distribution toward longer intervals (IPSCs: N=400 control events, N=397 baclofen events, p=1.12×10-10, Kolmogorov-Smirnov test; EPSCs: N=250 control events, N=204 baclofen events, p=4.54×10-4, Kolmogorov-Smirnov test). Summary of sIPSC (C) and sEPSC (D) statistics: (*i*) mean frequency, (*ii*) peak amplitude, (*iii*)10-90 rise time, and (*iv*) single exponential decay time constant before (Con) and after baclofen application (Bac). A significant decline in mean event frequency was observed for sIPSCs (n=20 experiments, *** p=1.9×10^−6^, Wilcoxon rank sum test) and sEPSCs (n=9 experiments, * p=0.00122). No significant differences were found in sIPSC amplitude, rise time or decay kinetics following baclofen application (sIPSCs, n=6; sEPSCs, n=6).

Likewise, cumulative histograms of the interevent intervals of sEPSCs showed significant shifts to longer intervals (N=250 control events, N=204 baclofen events, p=4.54×10^−4^, Kolmogorov-Smirnov test, Fig. 3Bii). The average spontaneous event rate sEPSCs was reduced from 1.5±1.3 Hz in control conditions to 1.0±1.9 Hz in baclofen (n=9, p=0.012, Wilcoxon rank sum test, Fig. 3Di). In contrast, baclofen application did not significantly affect the amplitudes and kinetics of either the sIPSCs (Fig. 3Cii-iv) or sEPSCs (Dii-iv). Spontaneous IPSCs had a mean amplitude of 0.11±0.56 nA in control and 0.091±0.059 pA in baclofen (n= 6, p=0.59), while sEPSCs had a mean amplitude of 0.040±0.015 nA in control and 49.4±0.018 nA in baclofen (p=0.44). Spontaneous IPSCs had single-exponential decay time constants of 12.6±7.5 ms in control and 19.1±15.9 ms in baclofen (p=0.56), while sEPSCs had decay time constants of 0.96±0.61 ms in control and 0.99±0.85 ms in baclofen (p=0.92). These data suggest that baclofen may act via presynaptic GABA_B_ receptors to alter release, but had little effect on postsynaptic determinants of the neurotransmitter response.

### Effects of GABA_B_R-mediated suppression on plasticity

To determine how GABA_B_ receptor activation might affect synaptic dynamics, we measured the IPSC amplitude profile during partial baclofen suppression on 20 Hz trains (Figure 4A). In the majority of recordings, the responses to trains of stimuli at this frequency revealed transient or sustained facilitation of the IPSC amplitude under control conditions (Fig. 4B, black trace). Because the maximal baclofen application (20 μM) could sometimes reduce the initial response amplitudes to negligible levels (e.g. ‘max baclofen’, Fig. 4A,B), changes in short-term plasticity were evaluated using blocks of trials during which the initial IPSC was reduced to 38% of the control amplitude (Fig.4A-D)(control: −0.29±0.16 nA; ‘partial baclofen’: −0.11±0.08 nA, n=7, p=0.016). As baclofen reduced the initial IPSC amplitude, the relative amplitudes of IPSCs later in the train were enhanced, showing greater facilitation compared to control (Fig. 4B,C; p=1.2×10^−5^, 2-way ANOVA). While the paired pulse ratio marginally increased from 1.28±0.46 to 1.58±0.88 (22% increase, p=0.43, no significant difference), the steady state ratio doubled from 1.24±0.64 to 2.47±1.63 (99% increase, p=0.0495)(Fig. 4Ei,ii).

**Figure 4.**
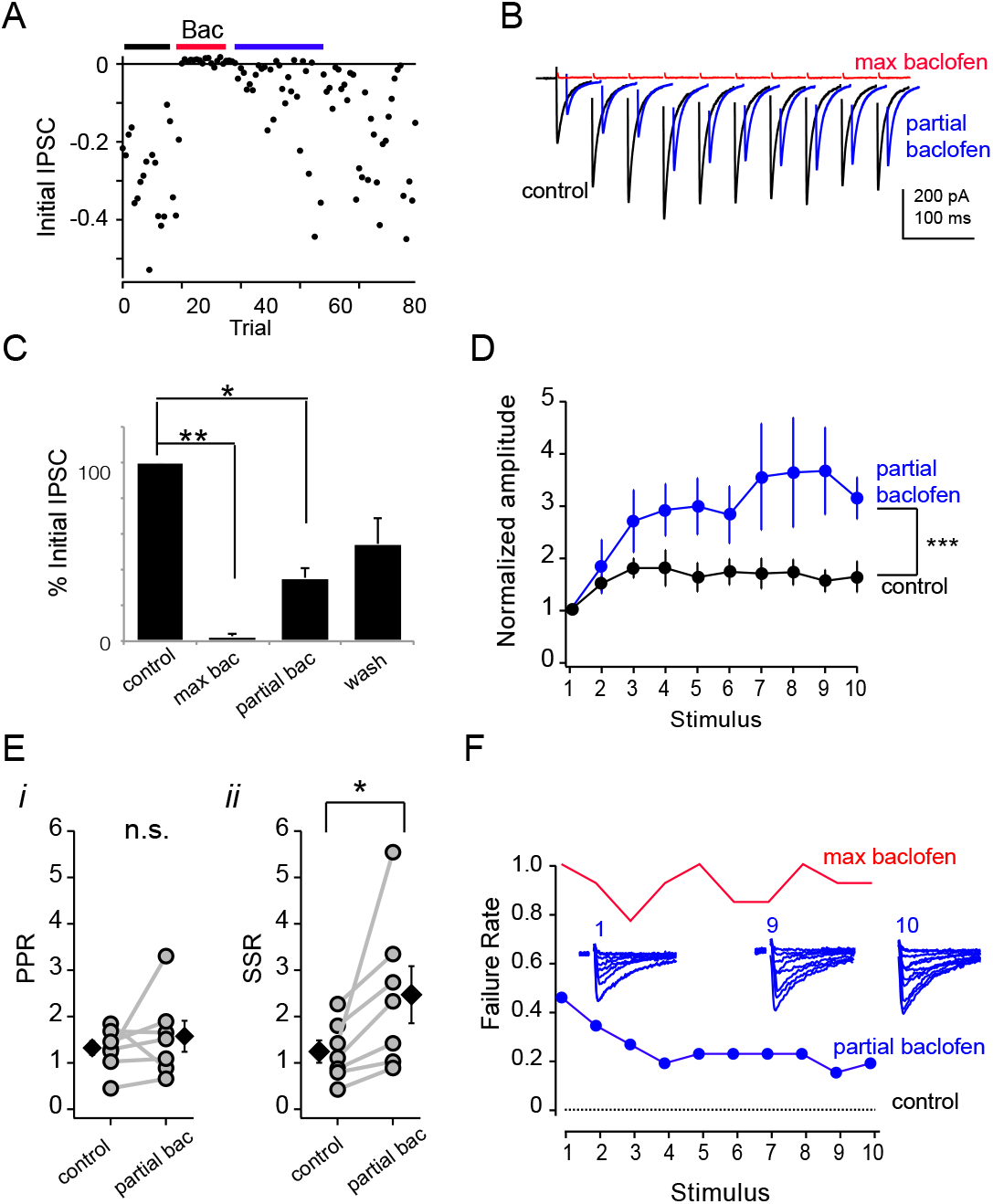
Effect of baclofen on short-term synaptic plasticity during trains of electrically evoked IPSCs. Plasticity of IPSC train responses assessed during periods of partial GABA-B receptor activation with baclofen revealed enhanced facilitation. A) Amplitude of the initial evoked IPSC of the train was profoundly suppressed during drug application (red bar), then recovered gradually. Blue bar shows time frame of trials analyzed for partial recovery. B) Comparison of mean current traces of train (20 Hz) before (black) and after maximal suppression (red), and during partial recovery (blue traces are horizontally offset for clarity). C) Normalized amplitude plots shows short-term plasticity of IPSCs over train. Sustained facilitation was typical of IPSC dynamics during moderate stimulation frequencies during control trials (black). Baclofen suppressed initial IPSC amplitude, but enhanced relative facilitation (blue). Mean±s.e.m. for 7 experiments; *, significant difference using Bonferroni adjusted pairwise Student’s paired t Test. D) Initial PSC suppression during control, maximal blockade (>90% blocked, **p<0.01) partial blockade (>64% blocked, * p<0.05) and late washout (not significantly different from control). E) Paired pulse ratio (PPR) during trains was unchanged during partial baclofen blockade (*i*), but sustained facilitation of later IPSCs in the train (SSR, steady state ratio) increased significantly (*, p<0.05)(*ii*). F) Failure rate at each stimulus in the train, uner control conditions (black line~0), partial blockade (blue markers and line), and maximal blockade (red line). Partial baclofen suppression revealed decreasing failure rate over the course of the train. Inset traces show responses during partial blockade at the first (1), ninth (9), and tenth (10) stimulus of the train.

Partial blockade allowed investigation of how release probability changed over the course of the train by measuring the failure rate. Under control conditions, the evoked IPSC failure rate in response to the first stimulus was nearly zero (1.3±1.9%, n=7 experiments); as these were not minimal stimulation experiments, responses likely comprised multiple inputs. In contrast, during partial blockade the failure rate in response to the first stimulus was ~40% (one example shown in Fig. 4F, ‘partial baclofen’, blue line). Over the course of the train, the failure rate then declined to ~19% in response to the last stimulus. Over 7 experiments, during partial blockade the failure rate was 38.8±31.3% for the initial IPSC and declined to 17.2±0.3% by the end of the 20 Hz train. During maximal blockade, we could also detect some decline in the failure rate over the train (Fig. 4F, ‘max baclofen’, red line). For all experiments, under maximal blockade, the failure rate in response to the first stimulus was on average 80% (79.6±21.6%, n=7), which then decreased over the course of the train to 50% (49.9±37.9%). Together with the effects observed on the spontaneous neurotransmitter release, these results suggest that activation of GABA_B_ receptors acts by reducing the initial release probability, which then increased over the course of the train via enhanced presynaptic facilitation.

Similar experiments were performed to determine how GABA_B_ receptor activation affected the excitatory synaptic dynamics. In these experiments, under control conditions the EPSC amplitude depressed to approximately 60% of initial amplitude during a 20 Hz constant frequency train (Fig. 5A, B; control: black trace, partial baclofen: blue trace). To measure whether baclofen had comparable effects as on IPSCs, short-term plasticity of the EPSCs was evaluated using blocks of trials during which the initial EPSC was reduced approximately to ~26% of the control amplitude (control: 1.46±0.73 nA; ‘partial baclofen’: −0.39±0.18 nA, 74% reduction, n=5 neurons). As baclofen reduced the initial EPSC amplitude, the relative amplitudes of EPSCs later in the train were enhanced, showing less depression compared to control (p=0.00412, 2-way ANOVA). Under partial baclofen suppression, the paired pulse ratio for EPSCs showed no significant change (control: 0.80±0.17, baclofen: 0.92±0.12, n=5, p=0.24, Student’s paired t Test)(Fig. 5Di), while the steady state ratio increased significantly by 44% (control: 0.52±0.10, baclfoen: 0.75±0.17, p=0.031). Thus under conditions of a moderate, but not maximal, activation of GABA_B_ receptors, there was a small but significant shift in short-term plasticity towards less depression in their excitatory train responses.

**Figure 5.**
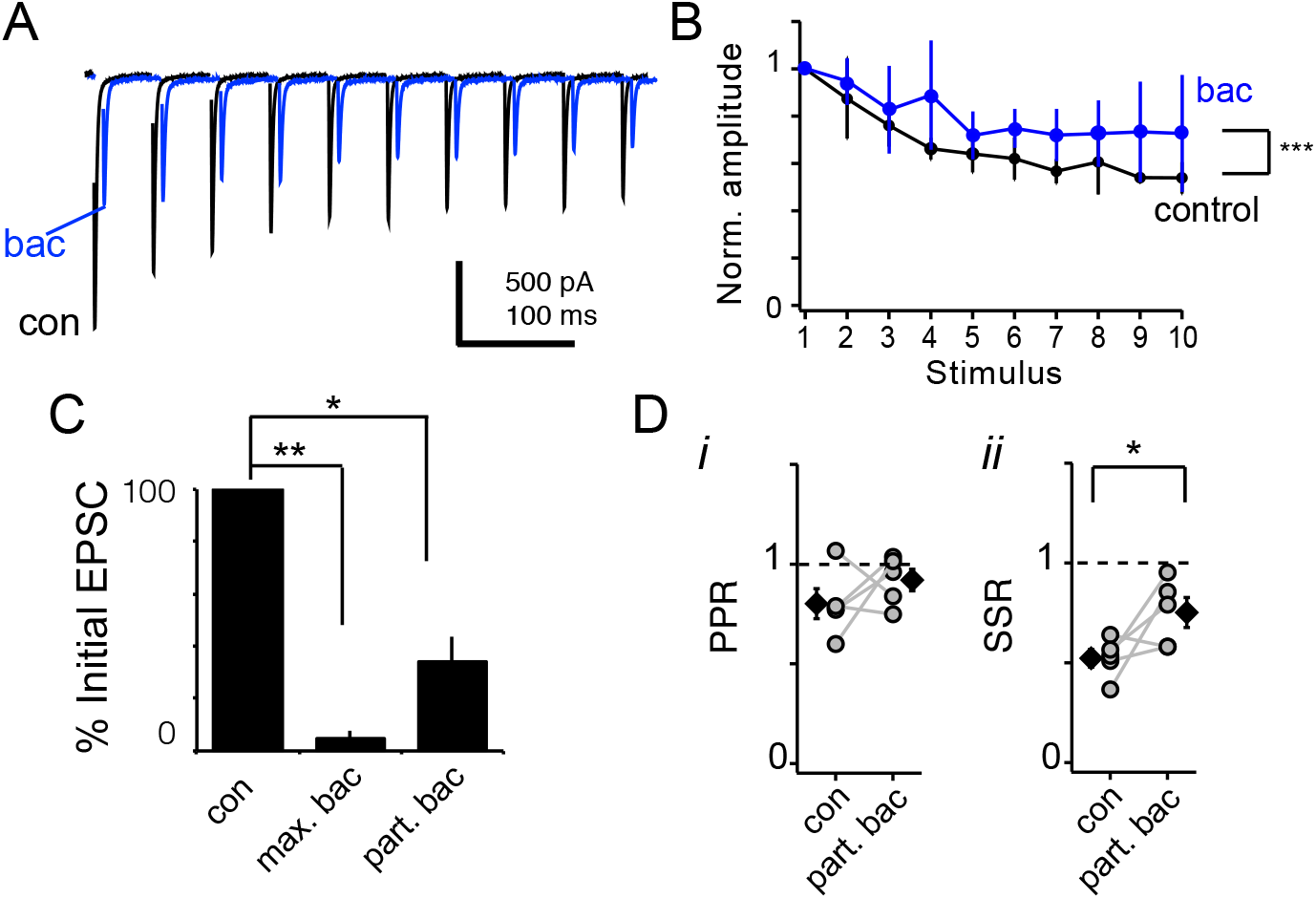
Partial GABA-B receptor activation with baclofen altered the short-term synaptic plasticity of EPSCs during short trains. A) Comparison of mean traces of train of EPSCs (20 Hz) before (black trace) and after partial suppression with baclofen (blue trace) in one recording (traces are horizontally offset for clarity). B) Normalized amplitude plots shows short-term plastitiy of EPSCs over train. Baclofen suppressed initial EPSC amplitude, but enhanced relative amplitudes (blue). Mean±s.e.m. for 4 experiments; posthoc testing did not reveal significant differences. C) Initial EPSC suppression during control, maximal blockade (>90% blocked, **p<0.01), and partial blockade (>66% blocked, * p<0.05) showed comparable levels to IPSC experiments. D) Paired pulse ratio (PPR) of the EPSCs during trains was unchanged during partial baclofen blockade (*i*), but the short-term synaptic depression of later responses in the train decreased significantly (steady state ratio increased, SSR, *, p<0.05) (*ii*).

### Evidence for basal GABA_B_ receptor activity on synaptic weight

To test whether GABA_B_ receptors could endogenously modulate synaptic dynamics during physiological activation in vitro, we elicited trains of evoked IPSCs before and after applying an antagonist of the receptor, CGP52432 (10 μM), and measured the effects on short-term plasticity. Unexpectedly, bath application of CGP52432 increased the initial IPSC amplitude (Fig. 6A). On average, the initial IPSC increased by 41% (control: −0.44±0.29 nA, CGP52432: −0.62±0.38 nA, n=10, p=0.018, paired Students *t*-Test)(Fig. 6Ai). Conversely, inhibiting GABAergic reuptake with NNC771 (20 μM) decreased the evoked IPSC amplitude by 30% (control: −0.51±0.26 nA, NNC771: −0.36±0.22, n=10, p=0.022)(Fig. 6Aii). No significant effects were observed by either drug on the amplitude of evoked EPSCs (initial EPSC NNC771 experiments: control −0.84±0.17 nA, NNC, −0.73±0.28 nA, n=4, p=0.31; initial EPSC CGP52432 experiments: control −1.08±0.72 nA, CGP52432 −1.05±0.42 nA, n=6, p=0.83)(Fig. 6B).

**Figure 6.**
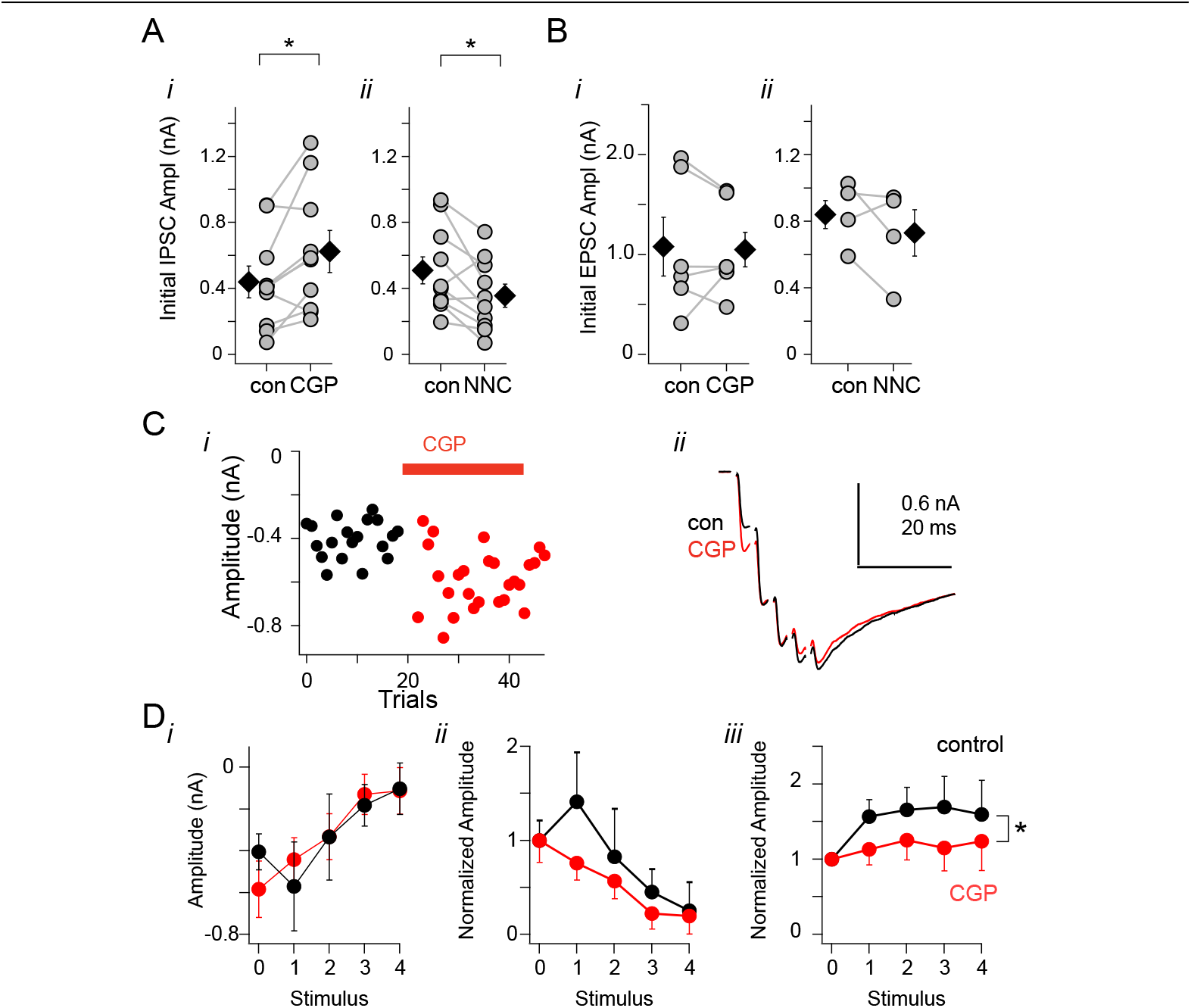
Blocking GABA reuptake (with GABA trasnsporter inhibitor NNC 771) or GABA_B_ receptor (with antagonist CGP 52432) has opposing effects on baseline inhibitory synaptic release and short-term synaptic plasticity. A) Evoked IPSC amplitudes were enhanced with application of GABA_B_ receptor antagonist CGP 52432 (*i*) and suppressed with inhibition of GABA reuptake transporter NNC771 (*ii*). B) Evoked EPSCs were unaffected by either drug. Diamond markers in panels A and B show mean±s.e.m. C) Example experiment of CGP 52432 effects on short, high frequency (200 Hz) trains of IPSCs. CGP 52432 increased the amplitude of initial IPSC in the train (*i*), but not later IPSCs (*ii*). D) Measured net amplitude (summation subtracted) of each IPSC in the train before (black, “con”) and after application of CGP 52432 (red, ‘CGP’). Increased amplitude of the initial response eliminated facilitation and resulted in further depression. Examples of single neurons responses showing raw (*i*) and normalized (*ii*) amplitudes and summary normalized short-term plasticity (*iii*)(n=5 experiments, 2-way ANOVA). Error bars are ±s.e.m.

The changes observed in short-term plasticity during IPSC trains were consistent with the change in initial amplitude: an increase the initial amplitude under CGP52432 was accompanied by a decrease in facilitation (n=5, p=2.14×10^−4^, 2-way ANOVA). Conversely, we asked whether the reduction in the initial inhibitory synaptic amplitude in the presence of the GABA reuptake inhibitor would be accompanied by an increase in facilitation. However, blocking the reuptake of GABA during trains with reuptake inhibitor NNC 771 (20 μM) was found to have minor and inconsistent effect on the short-term-plasticity with no significant effect on the train facilitation overall (n=4, p=0.083, 2-way ANOVA).

The effects of CGP 52432 and NNC 771 on the basal release properties during evoked trains implied ongoing, baseline activation of GABA_B_R, suggesting that the GABA chronically in the milieu was sufficient to affect these receptors. To determine whether spontaneous release was also affected by the same pharmacological manipulations, we measured spontaneous IPSC frequency before and after application of NNC 771 (20-80μM; Fig. 7A-C) and CGP52432 (10-20 μM; Fig. 7D-F). When NNC 771 was applied, the distribution of the interevent intervals between sIPSCs showed a significant shift toward longer intervals (Fig. 7B)(N_control_=378, N_NNC_=324, p=3.2×10^−6^, Kolmogorov-Smirnov test), while the sIPSC frequency was reduced by 29% (1.9±1.4 Hz in control to 1.5±1.0 Hz in NNC771; n=8, p=0.107, Wilcoxon Rank Sum test)(Fig. 7Ci). In these neurons, NNC 771 application appeared to have small but detectable effects on the synaptic current kinetics and amplitude. The sIPSC amplitude showed a small but significant decrease (Fig. 7Cii; control, −86.5±37.9 pA; NNC, −67.0±36.4 pA, p=0.039, Wilcoxon Rank Sum test) and the decay time course slowed slightly (Fig. 7Civ; control, 7.3±3.6 ms; NNC 771, 10.0±5.9 ms, p=0.039, Wilcoxon Rank Sum test), but no change in the rise time was observed (Fig. 7Ciii; control, 1.3 ±0.2 ms; NNC, 1.5±0.3 ms, p=0.148, Wilcoxon Rank Sum test). These mild postsynaptic effects on quantal event amplitude and kinetics with NNC771 indicate possible desensitization of postsynaptic ionotropic GABA receptors (Tang and Lu, 2012b).

**Figure 7.**
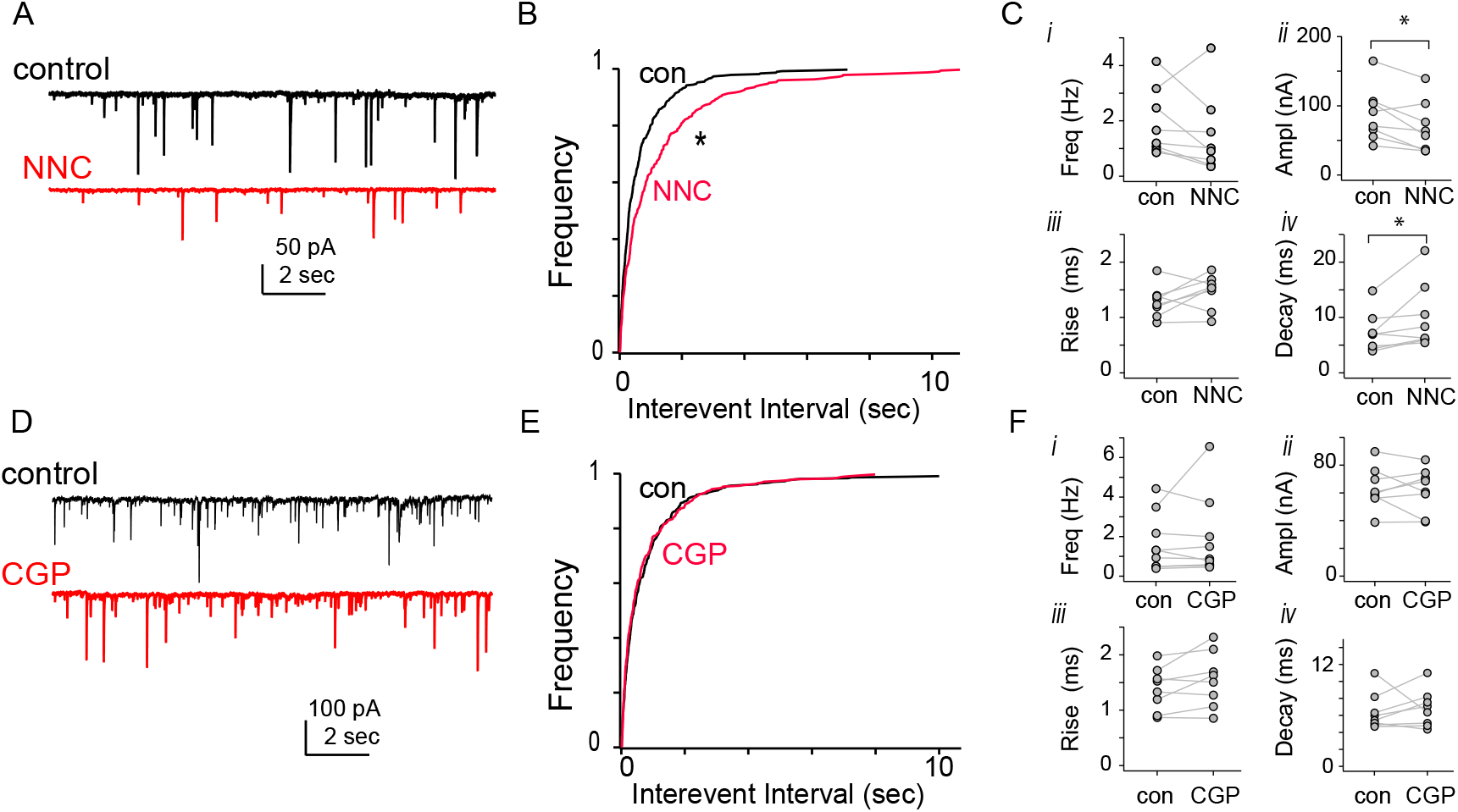
Effects of GABA reuptake inhibitor NNC771 (A-C) and GABA_B_ receptor blockade CGP 52432 (D-F) on spontaneous IPSC activity and kinetics. A) Example current traces showing spontaneous IPSC events before (top black trace) and after (bottom red trace) application of NNC 771. B) Cumulative histograms of sIPSC intervent intervals before and after application of NNC 771. The distribution of time intervals between events shifted toward longer intervals with drug application. C) Summary statistics on n=8 recordings showing mean sIPSC spontaneous rate frequency, peak amplitude, 10-90 rise time, and single exponential decay time constant before and after drug application. No signficant differences were found in average frequency or rise time; a signficant decrease in mean amplitude and increase in decay were observed (p=0.039 and p=0.039, Wilcoxon Rank Sum test). D) Example current traces showing spontaneous IPSC events before (top black trace) and after (bottom red trace) application of CGP 52432. E) Cumulative histogram of sIPSC intervent intervals before and after application of CGP 52432 showed no changes. F) Summary statistics reveal no signficant changes with CGP 52432 application showing mean sIPSC spontaneous rate frequency, peak amplitude, 10-90 rise time, and single exponential decay time constant before and after drug application (n=8).

In contrast, GABA_B_ receptor antagonist CGP 52432 had no effect on the spontaneous IPSC frequency, amplitude or kinetics (Fig. 7D). The cumulative histograms of the interevent intervals of sIPSCs were unchanged by application of CGP 52432 (Fig. 7E)(N_control_=363, N_NNC_=338, p=0.656, Kolmogorov-Smirnov test), while the frequency marginally increased from 1.8±1.3 Hz in control conditions to 2.1±0.9 Hz in CGP 52432 (no significant effect; Fig. 7Fi). No changes in the kinetics or amplitude of the sIPSCs were observed with the antagonist (Fig. 7Fii-iv).

## Discussion

These results suggest that both excitatory and inhibitory inputs may be modulated by the release of GABA via metabotropic receptors. The functional role of presynaptic GABA_B_ receptors in sensory processing at synapses in the NA is as yet undetermined, but in vitro heterosynaptic and homosynaptic effects of GABA were similar in the three important ascending auditory brain stem nuclei. Direct inhibitory modulation of NA neurons occurs via postsynaptic synapses and ionotropic GABA and glycine receptors, but the release of GABA may also indirectly influence information processing in these circuits via presynaptic mechanisms. Heterosynaptic effects on the auditory afferent inputs and homosynaptic effects on the inhibitory terminals that were equally suppressive in their actions, which could maintain the excitatory versus inhibitory balance while providing common negative feedback.

### Presynaptic GABAergic inhibition at inhibitory and excitatory synapses in NA

Physiological evidence suggested that the strong suppression of inhibitory and excitatory evoked currents occurred via a presynaptic mechanism. An immunohistochemical study previously showed GABA_B_ receptors were expressed in both avian cochlear nuclei and electron microscopy data further verified the presynaptic localization, at least within NM (Burger et al., 2005b). The clear decline in the frequency of spontaneous events that occurred with the GABA_B_ agonist supports a presynaptic locus of action, in agreement with a previous study (Shi and Lu, 2017). At the same time, spontaneous synaptic events showed no significant changes in event amplitude or time course with direct pharmacological activation of the GABA_B_ receptor, suggesting there was suggesting no postsynaptic effect on the channels that underlie the ionotropic IPSCs or EPSCs. Although evoked IPSCs showed a slightly faster time course, this could be explained by a reduction in asynchronous release that occurred in some recordings, a common feature with inhibitory inputs onto neurons inNM (Tang and Lu, 2012b) and onto spherical bushy cells in the AVCN (Nerlich et al., 2014). Although we did not further test presynaptic mechanisms, presynaptically localized GABA_B_ receptors have been shown in other systems to suppress neurotransmitter release via associated G-protein pathways and their downstream effectors. Downregulation of neurotransmitter release may occur by Gβγ subunit suppression of presynaptic voltage gated calcium channels or direct binding of vesicle release proteins inhibiting fusion (Gassmann and Bettler, 2012).

Previous research suggested that GABA_B_ receptor agonists could modify the excitability of NA neurons directly via postsynaptically located receptors (Shi and Lu, 2017). It is unlikely that the baclofen effects observed here are due to postsynaptic changes in membrane properties or voltage gated channel activation, however, as synaptic currents in this study were measured under voltage clamp conditions using a cesium intracellular solution expected to block potassium currents, one likely endpoint of postsynaptic GABA_B_ receptors. Even under current clamp conditions, baclofen did not induce changes in near-resting potential input resistance (Shi and Lu, 2017). Together these results suggest baclofen was acting solely via a presynaptic metabotropic pathway to modulate release in our study.

### Role of inhibitory feedback in the timing pathway

In the avian cochlear nucleus, the majority of known inhibitory feedback arises from an anatomically separate brainstem nucleus, the superior olivary nucleus (SON) (Funabiki et al., 1998; Yang et al., 1999; Lu and Trussell, 2000; Monsivais et al., 2000; Lu and Trussell, 2001). The two divisions of the cochlear nuclei, NA and NM, receive a common inhibitory feedback projection from the ipsilateral SON, as does the tertiary nucleus, NL (Burger et al., 2005a). NM and NL together comprise the circuit known for binaural processing of timing information important for localization sound in the azimuthal plane. How the inhibitory feedback affects the ascending sound information processing and whether there is functional heterogeneity in those effects across brain structures is yet to be fully understood. Because the timing circuit is highly specialized, while the ascending pathways via NA deal with sound intensity and spectrotemporal processing more broadly (Köppl and Carr, 2003; Steinberg and Peña, 2011; Fontaine et al., 2014), inhibition could have quite different roles in each area.

As sister nuclei that receive similar ascending nerve input and a common feedback, it is not surprising NA and NM have a number of similarities in their inhibitory synaptic properties. Among these are: co-release of GABA and glycine from presynaptic inhibitory terminals, likely arising from the same SON axons branching and projecting to each nucleus; chloride reversal potentials that are well above resting potential, resulting in depolarizing, inward currents; expression of postsynaptic fast ionotropic GABA_A_ receptors (GABA_A_R), postsynaptic glycine receptors (GlyR), and presynaptic and postsynaptic GABA_B_ receptors (Brenowitz et al., 1998; Burger et al., 2005b; Kuo et al., 2009; Fischl and Burger, 2014; Shi and Lu, 2017). However, there are also a number of differences whose functional importance have yet to be explained. For example, in NA, the ionotropic postsynaptic inhibitory currents are mixed GABA_A_R /GlyR currents, while in NM (and NL) the inhibitory currents are largely carried by GABA_A_R (but see Fischl et al., 2013 and Fischl and Burger, 2014 for a possible role by GlyR during high activity levels). Furthermore, the inhibitory currents measured in NA are faster than in NM and NL, likely due to the faster kinetics of the GlyR component (Kuo et al., 2009).

Our findings suggest that the modulation of excitatory and inhibitory inputs of NA neurons via GABA_B_R activation appears to mirror previous studies of GABAergic modulation of synaptic transmission in NM (Brenowitz et al., 1998; Brenowitz and Trussell, 2001a; 2001b). Agonists of GABA_B_R had an strong presynaptic suppressive effect on excitatory, presumptive nerve, inputs to NM neurons. These axosomatic calyceal terminals are highly specialized endings thought to reliably evoke large, suprathreshold glutamatergic potentials sufficient to elicit an action potential without fail. These relatively high-release probability synapses also show strong activity-dependent synaptic depression. Upon activation of presynaptic GABA_B_ receptors, the baseline release probability was reduced and the initial amplitude was reduced (but still suprathreshold), but later responses in a train of activity were, paradoxically, less likely to fail to produce action potentials due to reduction in synaptic depression and relief from receptor desensitization.

In NA, presynaptic modulatory effects may similarly alter the synaptic dynamics: GABA_B_R activation likewise suppressed baseline release and shifted the synaptic plasticity away from depression. However, because the excitatory inputs to NA neurons tend to be subthreshold (requiring summation to elicit an action potential) and exhibit less synaptic depression than NM under control conditions, presynaptic GABA_B_ modulation could have markedly different end effects on this network. Excitatory inputs to NL, which originate from NM, do not appear to be modulated by presynaptic GABA_B_ receptors, suggesting differential regulation of excitation by SON feedback relative to the cochlear nuclei (Lu et al., 2005; Tang et al., 2009).

### Heterosynaptic versus homosynaptic inhibition

In summary, we conclude that inhibitory terminals from the SON are subject to negative feedback or self-regulation in all three nuclei to which it projects. Thus the SON provides 1) direct inhibitory input to the projection neurons of NA via ionotropic GABA/glycine receptors (Kuo et al., 2009), 2) indirect, heterosynaptic inhibitory control of excitatory drive via metabotropic GABA_B_ receptors on glutamatergic terminals (Lu et al., 2005; Lu, 2009), and 3) a negative feedback regulatory signal via homosynaptic inhibitory metabotropic GABA_B_ receptors on the SON terminals (this study). The agonist effectiveness on the excitatory and inhibitory release appeared to be comparable, causing >90% suppression. Under what physiological conditions modulation by presynaptic GABA_B_ receptor occurs is yet to be determined, but our experiments suggest the inhibitory terminals may be more susceptible to modulation. During mild to moderate stimulation conditions, GABA release was sufficient to suppress baseline transmission at inhibitory synapses (see Fig. 6). In addition, blocking reuptake with an inhibitor of the GABA transporter, presumably causing transmitter pooling in the synaptic cleft, was sufficient to reduce spontaneous inhibitory event frequency, but not spontaneous excitatory event frequency (see Fig. 7). On the other hand, blocking GABA_B_ receptors, however, had no effect on spontaneous inhibitory (nor excitatory) event frequency, suggesting ambient spontaneous neurotransmitter release is not sufficient to elicit chronic, baseline suppression via this pathway. These results suggest that under normal physiological activity, the SON input may itself be under chronic, mildly suppressive negative feedback.

Our experiments failed to produce evidence for physiologically-evoked, endogenous heterosynaptic suppression of excitatory inputs to NA neurons. Neither NNC771 nor CGP52432 had any effect on the initial or singly-evoked EPSCs (see Fig. 6) or on the short-term plasticity during EPSC trains (data not shown). Because the EPSCs were pharmacologically isolated, it could not be determined whether SON fibers were also releasing GABA under the same stimulation paradigm that elicited the EPSCs. Nevertheless, our current results suggest that previous studies on synaptic plasticity using trains of stimulation in conditions in which metabotropic receptors were unblocked were probably not complicated by modulatory effects in slice (MacLeod et al., 2007). Heterosynaptic effects that rely on neurotransmitter spillover may be more likely in NM, where large calyceal axosomatic synapses interdigitate with small, punctate axosomatic inhibitory terminals. Whether the relative position of excitatory and inhibitory synaptic inputs to NA neurons facilitates such indirect modulatory effects in the intact circuit remains to be investigated.

## Acknowledgements

Support for this research was provided by National Institutes of Health NIDCD research grant R01DC10000 (K.M.M.). The authors thank P. Timi for assistance with analysis.

